# From Habitat Use to Social Behavior: Natural History of a Voiceless Poison Frog, *Dendrobates tinctorius*

**DOI:** 10.1101/515122

**Authors:** Bibiana Rojas, Andrius Pašukonis

## Abstract

Descriptive studies of natural history have always been a source of knowledge on which experimental work and scientific progress rely. Poison frogs are a well-studied group of small Neotropical frogs with diverse parental behaviors, distinct calls, and bright colors that warn predators about their toxicity; and a showcase of advances in fundamental biology through natural history observations. The dyeing poison frog, *Dendrobates tinctorius*, is emblematic of the Guianas region, widespread in the pet-trade, and increasingly popular in research. This species shows several unusual behaviors, such as the lack of advertisement calls and the aggregation around tree-fall gaps, which remain poorly described and understood. Here, we summarize our observations from a natural population of *D. tinctorius* in French Guiana collected over various field trips between 2009 and 2017; our aim is to provide groundwork for future fundamental and applied research spanning parental care, animal dispersal, disease spread, habitat use in relation to color patterns, and intra specific communication, to name a few. We report sex differences in habitat use and the striking invasion of tree-fall gaps; describe their courtship and aggressive behaviors; document egg development and tadpole transport; and discuss how the knowledge generated by this study could set the grounds for further research on the behavior, ecology, and conservation of this species.

## Introduction

Natural history has been long acknowledged as the foundation of new hypotheses in behavioral and evolutionary ecology (Endler 2015). Thus, scientific progress relies greatly on knowing what different organisms are, where they live, what they feed on, how they respond to different stimuli, and what kind of other peculiar behaviors they exhibit (Tewksbury et al. 2014). This, more often than not, is achieved through field observations.

Neotropical poison frogs (Dendrobatidae) and their close relatives are a showcase example of how detailed knowledge of natural history can lead to groundbreaking hypothesis-driven studies (e.g. Santos et al. 2003; Brown et al. 2010; Amézquita et al. 2011; Pašukonis et al. 2014; Tarvin et al. 2017). Exhaustive field observations have revealed the diversity of poison frog parental care behavior (e.g. Crump 1972; Silverstone 1973; Donnelly 1989; Summers 1989; Caldwell 1997), warning coloration (e.g. Silverstone 1975; Myers and Daly 1983), and skin alkaloids (e.g. Myers and Daly 1976, 1980; Brodie and Tumbarello 1978; Myers et al. 1978; Summers 1989), aspects that have become a trademark in the group both for research and for the pet-trade. However, there is still a surprising lack of information on the natural history of some species that have become increasingly well studied otherwise, such as the dyeing poison frog, *Dendrobates tinctorius*.

Although bred in captivity by hobbyists for decades (Schmidt and Henkel 1995; Lötters et al. 2007), and despite its growing status as a model species for studies on the evolution and function of coloration (e.g., Wollenberg et al. 2008; Noonan and Comeault 2009; Rojas et al. 2014a, 2014b; Barnett et al. 2018), there are only a handful of studies on *D. tinctorius* in its natural environment; most of these have been carried out and published only after 2010 (Born et al. 2010; Courtois et al. 2012; Rojas and Endler 2013; Rojas 2014, 2015; Rojas et al. 2014a). Four other studies in the wild have attempted to understand evolutionary aspects of their variable coloration, using clay or wax models instead of the actual frogs (Noonan and Comeault 2009; Comeault and Noonan 2011; Rojas et al. 2014b; Barnett et al. 2018).

Many poison frog field studies over the last five decades have relied on prominent male calls either directly, by studying aspects related to vocal behavior (e.g., Fandiño et al. 1997; Lötters et al. 2003; Forsman and Hagman 2006; Erdtmann and Amézquita 2009; Vargas-Salinas and Amézquita 2013), or indirectly, by using the calls to locate territorial males in the field (e.g., Bee 2003; Hödl et al. 2004; Amézquita et al. 2005; Rojas et al. 2006). Meanwhile,*D. tinctorius* remained almost unstudied, at least in part, due to their lack of a regular calling behavior. Therefore, much of the behavioral and evolutionary ecology of dyeing poison frogs remains unknown.

As stated by the IUCN Red List for Threatened Species, *D. tinctorius* is in the category ‘Least Concern’ (Gaucher and MacCulloch 2010). According to this report, its major threat is illegal trading, as it is for various other dendrobatid species (Gorzula 1996; Nijman and Shepherd 2010; Brown et al. 2011; Hoogmoed et al. 2012). However, a recent study provided evidence that, despite having seemingly large and stable populations throughout its range, *D. tinctorius* is not safe from the chytrid fungus (*Bd*) infection (which, incidentally, was discovered in a captive individual of *D. tinctorius;* Longcore et al. 1999) in its natural habitat (Courtois et al. 2012). Moreover, a recent study by Courtois et al. (2015) raised even greater concern as, of all the species tested for *Bd* in French Guiana, the highest prevalence was found in dendrobatid frogs, including *D. tinctorius*.

Alarming declines make it even more urgent to study the natural history of amphibian species and communities, especially of ‘sentinel’ species such as *D. tinctorius* (Courtois et al. 2015), whose declines provide anticipated warning of risks to human or ecosystem health (Beeby 2001). Only by understanding organisms in their own habitat can we produce sensible and timely conservation policies, and sustainable management (Tewksbury et al. 2014). In the particular case of *D. tinctorius*, knowing their habitat use, breeding biology, social behavior, and movement ecology could be of utmost importance for modeling disease spread and the impacts of deforestation, among other current environmental threats. Here, we document the habitat use, and the reproductive, social, and vocal behaviors of *D. tinctorius* in the wild; and provide information about various other aspects of its natural history that will be a valuable groundwork for future fundamental and applied research in behavior, ecology, evolution, and conservation.

## Materials & Methods

### Study species

*D. tinctorius* is a diurnal, relatively large (Snout-Vent Length 37—53mm at the study site; Rojas and Endler 2013) poison frog of the Neotropical family Dendrobatidae (more specifically of the *tinctorius* group; Grant et al. 2006), which occurs around canopy gaps in primary forests in the Eastern Guiana Shield, at elevations between 0 and 600 m (Noonan and Gaucher 2006; Wollenberg et al. 2006). It has skin alkaloids (Daly et al. 1987), and is characterized by a great color pattern variation both within (Fig. 1a; Rojas and Endler 2013) and among populations (Fig. 1b; Noonan and Gaucher 2006; Wollenberg et al. 2008).

**Figure 1.**
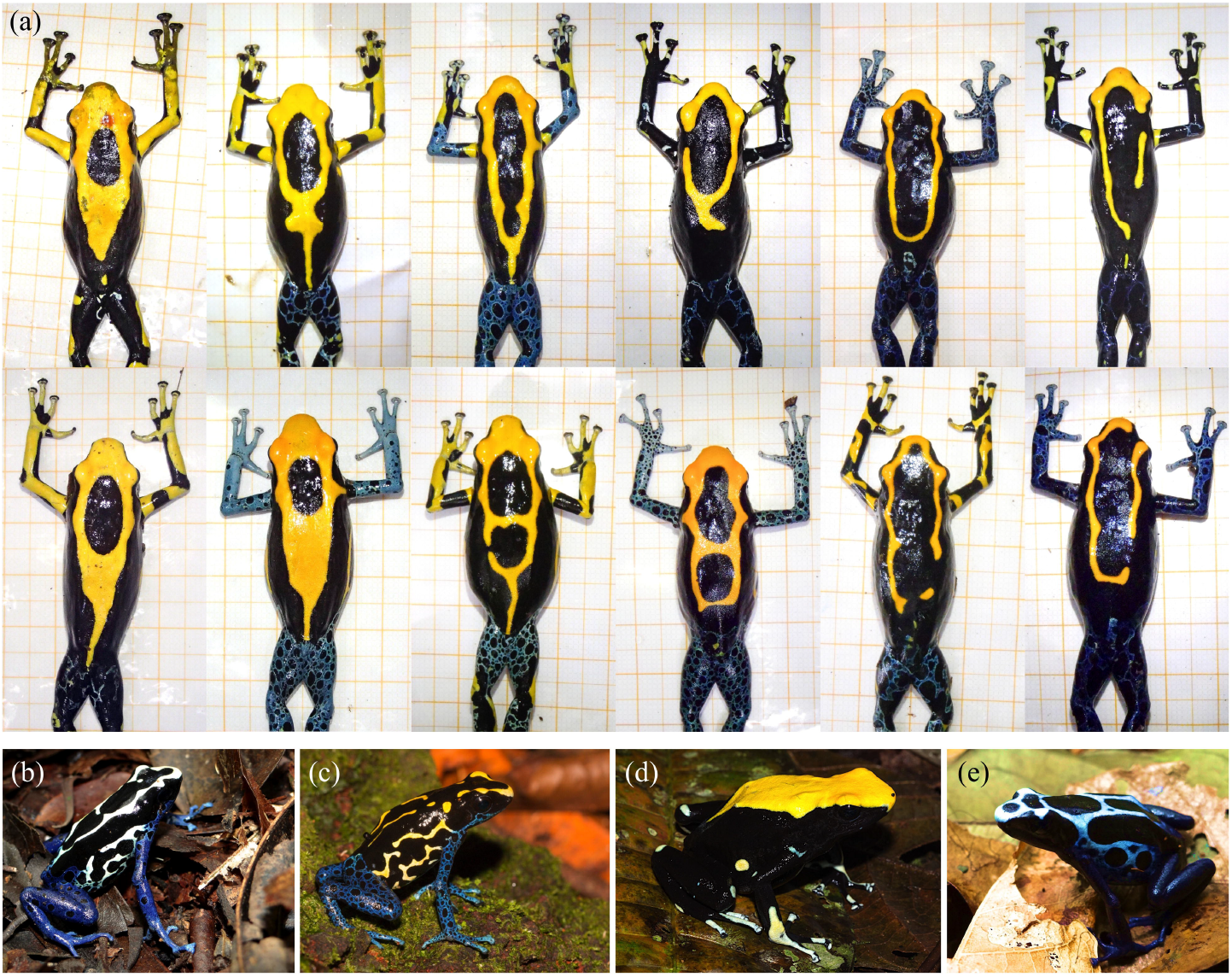
Colour pattern variation (a) within the studied population and (b-e) between different populations of the dyeing poison frog in French Guiana. (a) Top row males, bottom row females. Lines on the background paper mark 5 mm. Note the enlarged toe discs in males, but overall larger female body size. Photos by: A. Pašukonis and Matthias-Claudio Loretto (a: Nouragues nature reserve, French Guiana), A. Fouquet (c: Bakhuis, Suriname; d: Mt. Galbao, French Guiana), and B. Rojas (b: Mt. Matoury, French Guiana; e: Mt. Bruyere, French Guiana).

In our study area, color patterns can be used reliably for individual identification (Born et al. 2010; Courtois et al. 2012; Rojas and Endler 2013; Fig. 1a), and sex can be determined by the size of males’ toe discs, which are wider than females’ in relation to their body size (Rojas and Endler 2013). In contrast to most frogs (including closely related poison frogs), male *D. tinctorius* do not produce advertisement calls, and when they do vocalize, they do it very softly (Lescure and Marty 2000). Newly hatched tadpoles are carried by males to pools formed in tree-holes or palm bracts at variable heights (Fig.2; Supplementary video 1; Rojas 2014, 2015), where they remain unattended until metamorphosis, which occurs after approximately two months (BR, pers. obs. in the field). As in some other species of *Dendrobates* (Caldwell and De Araújo 1998, Gray et al. 2009, Summers 1990, Summers and McKeon 2004), larvae feed on detritus and on larvae of insects and frogs (BR, pers. obs.), including conspecifics (Rojas 2014, 2015; Supplementary movie 2). In captivity, individuals reach maturity after approximately 18 months (Lötters et al. 2007), but their age at sexual maturity in the field is unknown to date.

**Figure 2.**
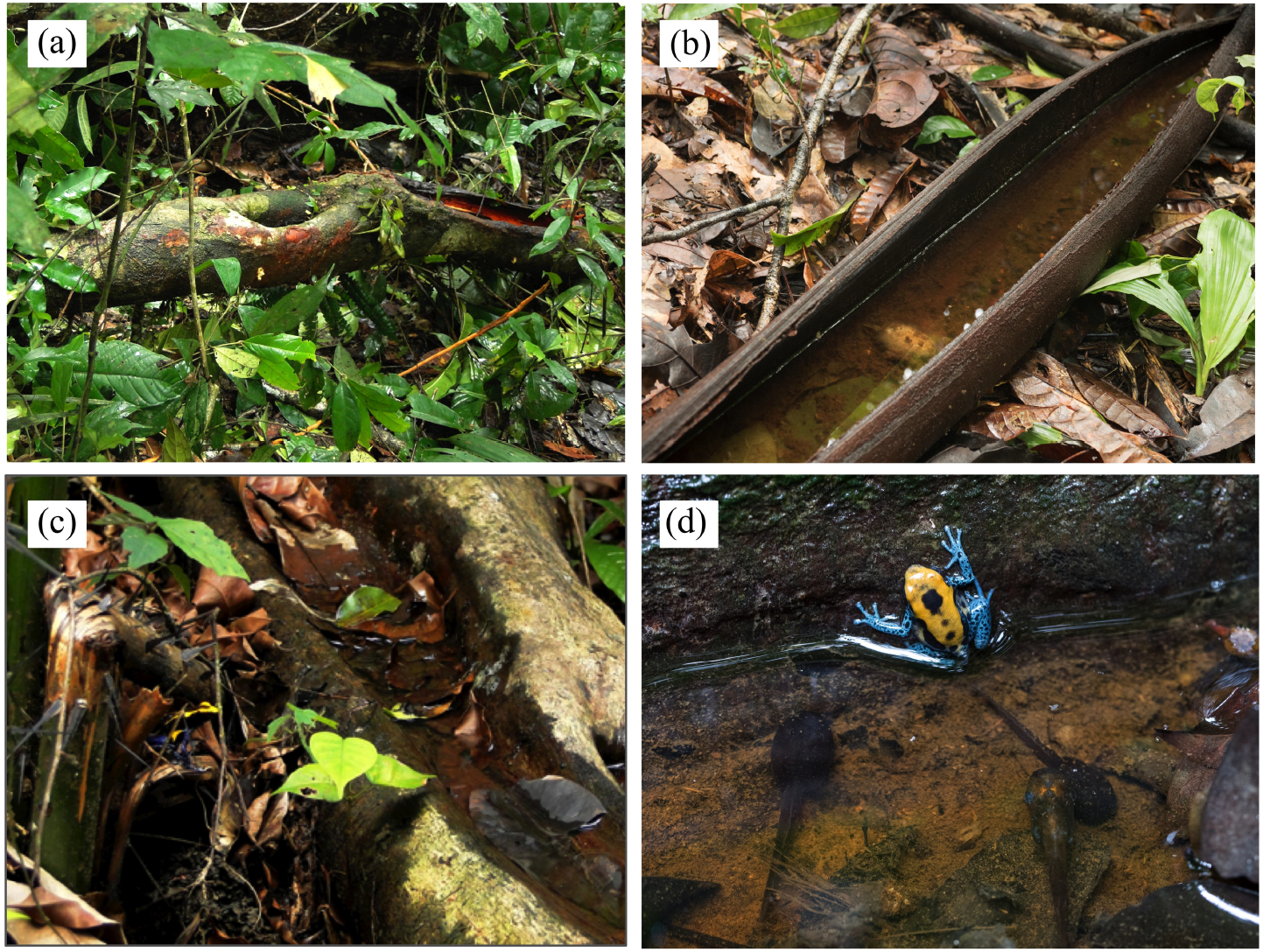
Examples of phytotelmata used as tadpole deposition sites at the studied population. Photos by: B. Rojas (a, c) and A. Pašukonis (b, d).

### Reproductive and social behavior

BR did systematic data collection during three field seasons between 9 January and 20 February 2009, 17 January and 19 March 2010, and 17 January and June 6 2011, at Camp Pararé, Nouragues Ecological Research Station, French Guiana (4°02’N, 52°41’W), in primary lowland terra-firme forest, where *D. tinctorius* is one of the most common leaf-litter frogs (Courtois, et al. 2013). In addition, AP made opportunistic observations on social and reproductive behavior at the same study site between January and March 2016 and 2017. The study periods correspond to the early rainy season and high reproductive activity of *D. tinctorius* in the study area.

During each study period between 2009 and 2011, BR surveyed a 1.5 km transect on a near-daily basis, between 8:00 and 17:30. Each frog found was captured, when possible, and photographed for future individual identification on the basis of its color patterns. When two individuals seemed to be interacting, they were followed for as long as it was necessary to determine the nature of the interaction (i.e. courtship or agonistic encounter). Two individuals were considered to be in courtship when they were less than 1 m apart (as in Pröhl 2002) and one was clearly following and touching the other (pers. obs.) for at least 15 min. A 15 min waiting time was chosen on the basis of previous studies of mate choice and assortative mating in captive dendrobatids (Maan and Cummings 2008, 2009). When possible, we followed pairs in courtship until they were no longer visible or until oviposition occurred, which proved difficult most of the time due to lack of accessibility and poor visibility under forest structures. Agonistic encounters were more difficult to follow because of their usually short duration and the high movement speed of the frogs, but we observed them for as long as both individuals were visible. Fragments of the two types of interactions were filmed for documentation purposes. Observations were done at irregular time intervals during the day.

Males carrying tadpoles were found during daily surveys along a 1.5 km transect. BR recorded the number of tadpoles on the back of each tadpole-carrying male and captured it when possible. Upon capture, each male was photographed (with the tadpole(s) still attached) against graph paper. Later these photos were used to measure the size of both the frog and the tadpoles with the software ImageJ (Schneider et al. 2012). Tadpole size was measured dorsally, from the tip of the snout to the base of the tail. The sizes of tadpoles carried by one male at the same time were compared (always the small vs. the large of each pair) with a paired *t*-test.

### Vocal behavior

*Dendrobates tinctorius* vocalizes rarely and at very low intensities, making it difficult to obtain audio recordings. We were able to obtain a high-quality audio recording of one male. In addition, to measure the acoustic properties of the call, we extracted lower quality audio from video recordings of social interactions. In total, we obtained sufficient quality recordings of eight calls produced by three males (four, three, and one call per individual). We manually measured the duration, pulse rate and dominant frequency of each call using Praat (v. 5.3.85; Boersma and Weenink 2014) acoustic analysis software. We averaged the measures between calls within each male and then between the three males. We used one call of the highest quality to visually illustrate call structure.

### Treefall-gap invasion

In a previous study, Born et al. (2009) reported frequent sightings of adult *D. tinctorius* in recently formed tree-fall gaps. BR witnessed the formation of nine tree-fall gaps over the study periods (one in 2009, eight in 2011); these were discovered rapidly because they occurred in the 1.5 km transect surveyed daily. BR inspected each gap within the first 24 hours of its formation and caught as many frogs as possible, moving fallen branches until no frogs were seen (after 2-3 hours). During the next two consecutive days BR carefully looked for new frogs over a similar period of time (2-3 h); most of the individuals seen then had been found on the first day. When frogs were seen but not caught, BR photographed them from a distance to record their color pattern for further identification upon capture. The presence of individuals in these tree-fall gaps continued to be recorded during daily surveys for up to two months after their occurrence.

### Habitat use

During the field season of 2010, BR captured 109 frogs (55 females and 54 males), each of which was assigned to one of two microhabitats according to where they were first seen: leaf litter (when frogs were on a relatively open patch of leaf litter without any obvious structure in a 1 m radius), or associated to the following structures: fallen logs (when frogs were visibly exposed on top of the log), fallen branches (when individuals were in fallen tree crowns) and tree/palm roots (when the frogs were within the exposed roots or next to them, or inside hollow trunks). Frogs were only included in the analyses once (recaptures of the same individual were excluded in order to avoid pseudoreplication, and only the site at first sighting was taken into account). We tested for differences between the sexes in the microhabitat where they were found (open vs. associated with the aforementioned structures) using a Generalized Linear Model with binomial distribution. All statistical analyses were done with the software R v. 3.3.3 (RCoreTeam 2014) using the RStudio interface (RStudio Team 2015).

### Ethics statement

Our research was approved and authorized by the scientific committee of the Nouragues Ecological Research Station. We strictly adhered to the current French and European Union law, and followed the Association for the Study of Animal Behaviour’s (ASAB) Guidelines for the use of live animals in teaching and research.

## Results

During three field seasons between 2009 and 2011, we identified 629 individuals unequivocally, 597 of which were captured. We photographed the remaining 32 frogs from a distance that allowed the record of their unique color patterns and, thus, their individual identification. There was no statistically significant difference between the number of females (N=276) and the number of males (n=321) found, although there was a non-significant trend towards a larger number of males (*χ*^2^= 3.392, df = 1, *P* = 0.066).

### Habitat use

We found clear differences between the sexes in terms of the microhabitat where they were found. Females were predominantly found in open areas of leaf litter (60% of females vs. 31.5% of males), whereas males were mostly found associated to structures (68,5% of males vs. 40% of females; estimate ± SE = 1.183 ± 0.402, *Z* = 2.943, *P* < 0.001, n=109; Fig. 3), specially fallen logs and branches (31.5% and 22.2%, respectively).

**Figure 3.**
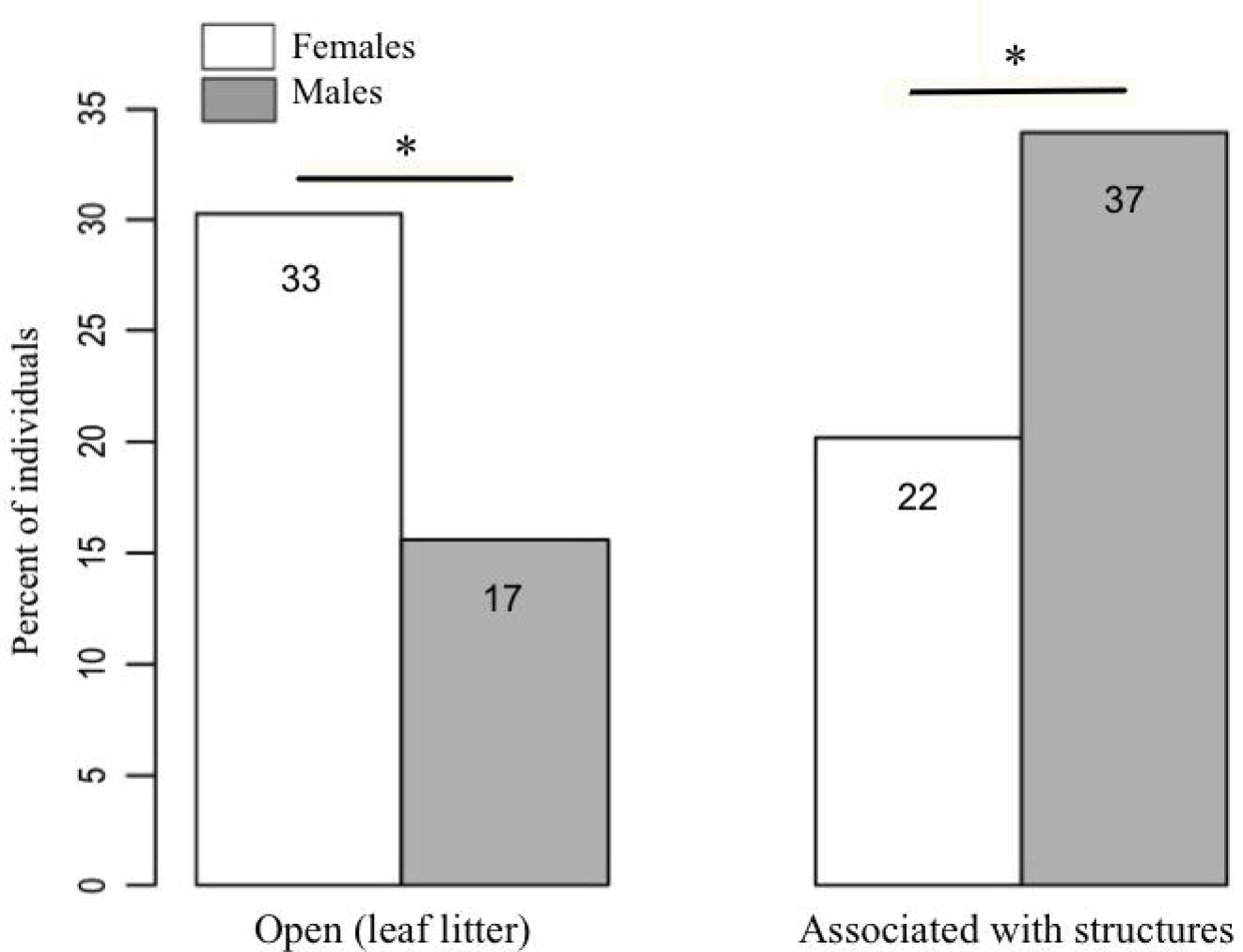
Habitat use in relation to sex. Numbers in the boxes indicate the total number of individuals in each category (N =109).

### Invasion of treefall gaps

A total of 140 individuals (65 females and 75 males) arrived in the nine fresh gaps studied, mostly on the same day of their occurrence or within the first three days. Males were as likely as females to arrive within this timespan (males: mean = 1.24 ± (SE) 0.08 d; females: mean = 1.08 ± (SE) 0.06 d; *χ*^2^= 0.714 df =1, *P* = 0.398; Fig. 4A). In the long-term (i.e., up to 60 days after the occurrence of the treefall), however, more males than females were found in treefall gaps (*χ*^2^= 11.137, df = 1, *P* = 0.001; Fig. 4B).

**Figure 4.**
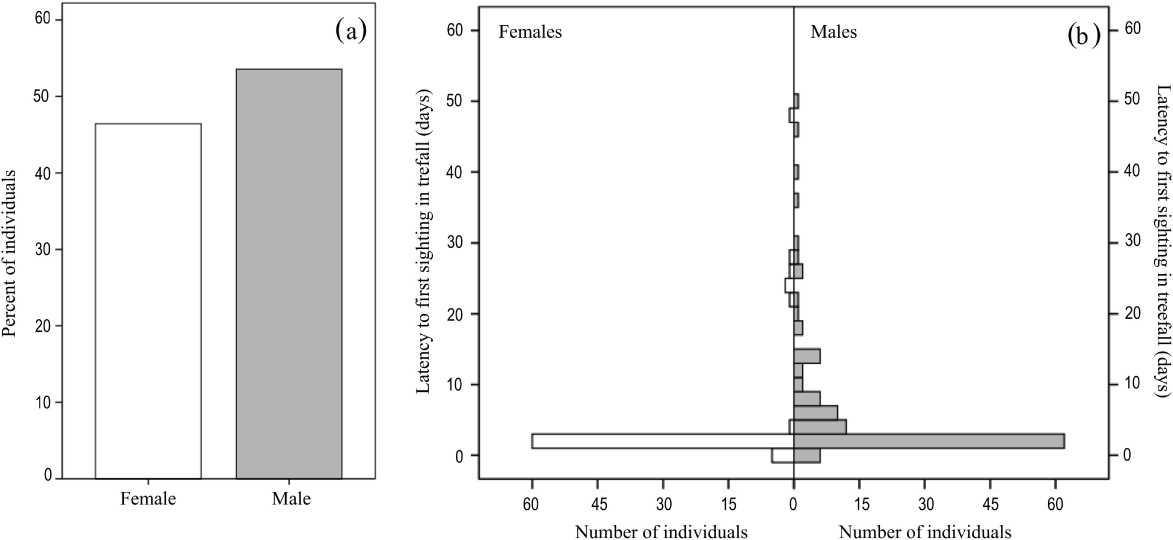
Dozens of adult *D. tinctorius* can aggregate at once at a newly formed treefall gap. There are no sex differences in arrival within the first three days of gap formation (a), but males are more likely to be found in treefall gaps in the long term (b).

### Vocal behavior

*Dendrobates tinctorius* produces a call that can be described as a very low intensity ‘buzz’ (Supplementary sound file). The call is audible to humans only from within a few meters; at times males inflate the vocal sac without anything audible to us from a distance of up to 1 m. Males call rarely and only when in courtship or during agonistic interactions with other males. We never observed a male calling alone. We were able to record and measure calls from two males in courtship and one in an agonistic interaction. Calls produced in courtship and agonistic contexts sounded similar to us and had similar acoustic parameters, although more recordings would be needed for a detailed comparison. All measured calls shared the same general structure: a short broadband burst of pulses produced at a high rate (Fig. 5B, C). The measured call duration was 0.55–0.98 s (mean = 0.76 s), the within-call pulse rate was 143–175 Hz (mean = 160 Hz), and the dominant frequency band centered around 2700–3270 Hz (mean = 3109 Hz).

**Figure 5.**
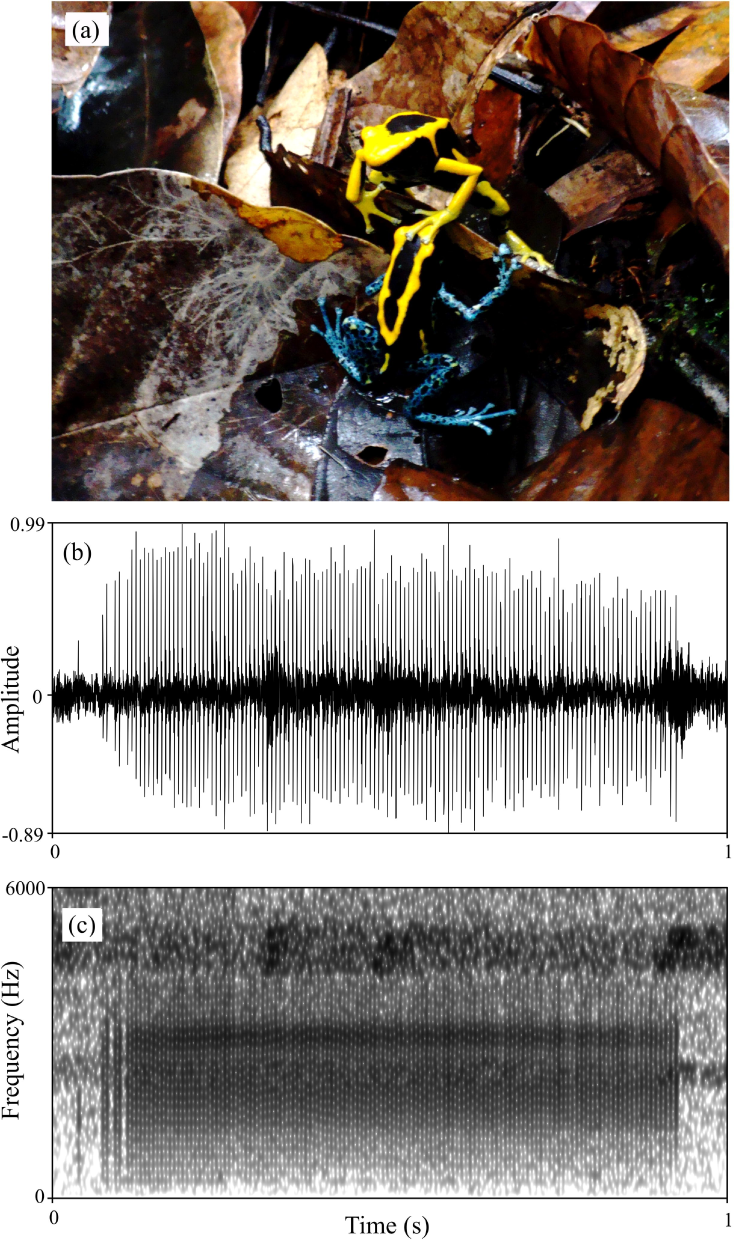
(a) Example of tactile interactions observed in *D. tinctorius* during courtship: a female with a limb on a male’s head. (b) Waveform and (c) spectrogram of *D. tinctorius* call recorded from close range (approx. 30 cm) during courtship. The normalised waveform reveals the relative amplitude modulation and the pulsating structure of the call (pulse rate = 154 Hz); the spectrogram (FFT window length = 0.01 s, Gaussian window, frequency range 0 - 6000 Hz) show the broadband spectral structure of the call with dominant frequency band centered around 3150 Hz. Photo by: B. Rojas.

### Courtship and egg laying

We found 47 pairs engaged in courtship (10 in 2009, 14 in 2010 and 23 in 2011), involving 40 males and 39 females. Courtship was observed throughout the day and lasted several hours. In one case a courting pair was followed for nearly 7 hours before oviposition took place. Courtship consists of several bouts of moving together and stationary tactile interactions (Fig. 5A, Supplementary video 3) that are interrupted, for example, when one of the individuals starts to feed. In general, each bout is initiated by tactile interactions in which the female repeatedly places one of her forelimbs on the male’s limbs, back or head. The male then usually faces her before moving away, followed by the female, in search of an egg-laying site. When a female stops following for several minutes, for example because she starts to forage, the male usually turns back and calls. Males also produce the same soft calls and vocal sac movements during some tactile interactions and following bouts. If the female does not approach the male, he occasionally approaches her and touches her head or back. Altogether, the courtship sequence in *D. tinctorius* appears to be very similar to that in *D. auratus* (Wells 1978), with females taking the most active role. Both males and females vibrate the second digit of the hind-legs at high frequency (‘toe trembling’, *sensu* Hödl and Amézquita 2001; see Supplementary video 3) during courtship. Toe-trembling behavior can also be observed during foraging and agonistic interactions.

The courting pair does not seem to move over great linear distances (mean = 4.5 m; range, 0–8 m; n = 6), but moves in circles within an area of a few square meters instead. Every now and then the pair stops at certain places under the leaves or inside a hollow trunk, and the female starts to move in circles on the same spot, with alternating movements of her hind limbs in what appears as wiping of the leaves (See Supplementary video 4). The pair sometimes rests on the same spot for several minutes and the tactile interactions increase considerably during these breaks. More often than not, the pair does this a few times, at different places, before they choose the place where egg laying occurs.

In addition to the clutches laid by pairs we followed during courtship (n=3), we found 18 clutches (for a total of 21) with 2–5 embryos (mean = 3.6) at different developmental stages. The eggs were laid under or within fallen logs and other wooden structures, leaf-litter, palm bracts and leaves, and animal burrows, usually completely sheltered from the rain (Fig 6A, B). Egg diameter is ~ 4.2 mm and hatching occurs after approximately two weeks (BR, pers. obs.; Fig. 6A). Eleven clutches were followed during development and only 14 out of 46 embryos (30,4 %) from eight out of 11 clutches survived until hatching. Other embryos did not develop, were destroyed by fungus, or disappeared possibly due to predation, although no predation was observed directly. Twelve clutches in total were observed with 1–4 (mean = 2.2) tadpoles ready for male transport (Fig. 6C). Males were found occasionally sitting near or on top of egg clutches, most likely inspecting and moistening them.

**Figure 6.**
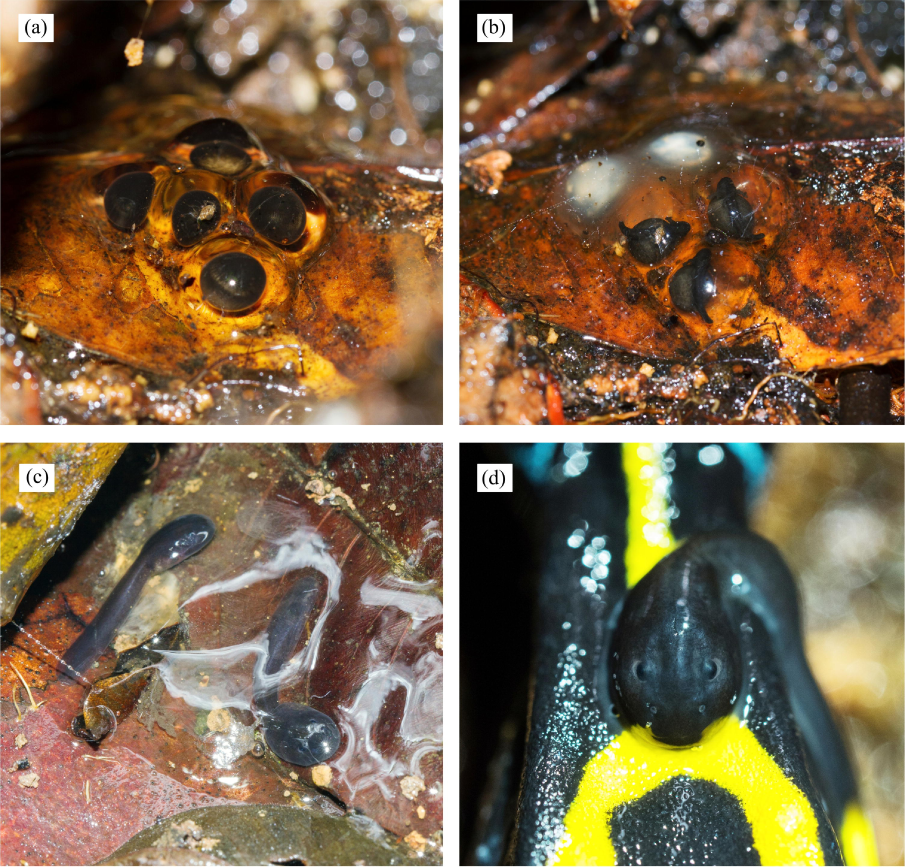
Clutch development in D. tinctorius in the field. (a) Freshly laid clutch of 5 eggs; (b) The same clutch five days later. Note that two of the initial eggs have been infected by fungus; (c) 15 days after egg laying, two surviving tadpoles are ready to be picked by the male and taken to a body of water where they will continue to develop until metamorphosis; (d) A tadpole attached to the male’s back. Photos by: A. Pašukonis.

### Larval development and patterns of tadpole transport

Hatching occurs after approximately 14 days (Fig. 6), but the tadpoles may remain viable in the clutch for several days before being transported (AP, pers. obs.). The male eventually returns and sits on the clutch, allowing the tadpoles to wriggle on his back, and takes them to suitable bodies of water where they will remain unattended until metamorphosis, feeding on detritus and the larvae of some insects (e.g., Diptera and Odonata) and other frogs (pers. obs.), even conspecifics (Fig. 7A; Rojas 2014, 2015). Tadpole mouthparts are well suited for their carnivorous diet, with hardened serrated jaw sheath (Silverstone 1975; Fig. 7B). Size at metamorphosis ranges 10.94 −15.62 mm (mean = 13.15 ± (SE) 0.24 mm, n=24), and at that point the color patterns are already completely visible (Fig. 2D).

**Figure 7.**
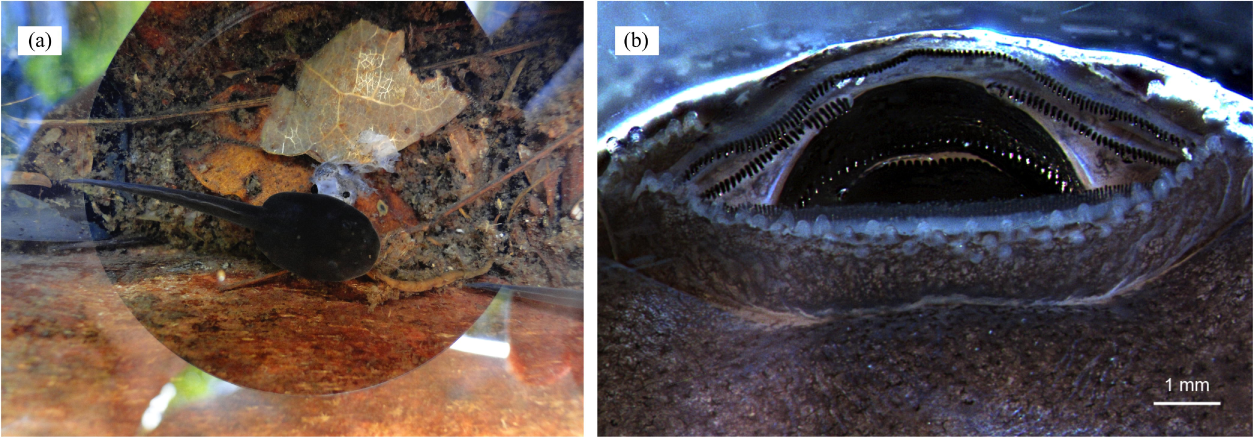
(a) A cannibalistic tadpole with the remainings of its victim; (b) oral apparatus (anterior side up) of a stage 25 (Gosner 1960) *D. tinctorius* tadpole. Photos by: B. Rojas (a) and E. K. Fischer (b).

We found 102 males (7 in 2009, 17 in 2010 and 78 in 2011) carrying one (Fig. 8A 79.4%), two (18.6%) or three (2.0%) tadpoles (mean = 1.23 ± (SD) 0.465; Fig. 8c) ranging 4.78–6.87 mm long (from the tip of the snout to the base of the tail; mean = 5.52 ± (SD) 0.50 mm). On one exceptional occasion BR also found one female carrying two tadpoles with a visible difference in size (Fig. 8B). Pairs of tadpoles transported by a male simultaneously differed significantly in size (paired *t*-test: *t*= 4.719, df=13, n=14, *P* <0.01; Fig. 8D). The tadpoles of the two males carrying three at a time (n=2) were excluded from this analysis. Some males carrying more than one tadpole were seen depositing one of them in a pool and leaving with the second tadpole still attached to their back, whereas other males were seen depositing their two tadpoles in the same pool, at the same time. Some males were also seen visiting more than one pool before the tadpole(s) detached from their back. The visits consisted of jumping into the pool and sometimes repeatedly diving inside for several minutes while the tadpole remained attached (see Supplementary video 1).

**Figure 8.**
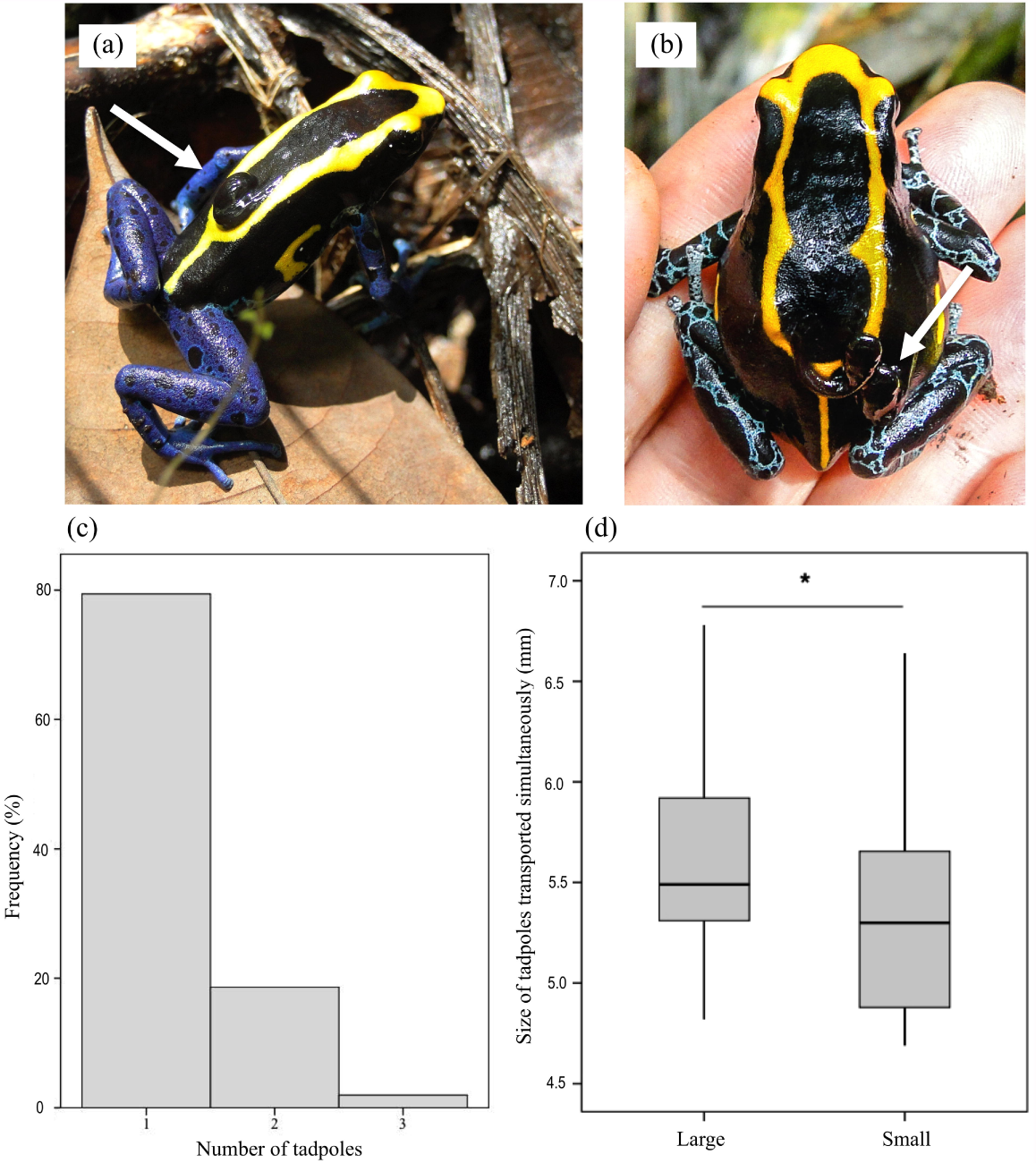
A male (a) and a (exceptional) female (b) with tadpoles on their back (indicated by white arrows). (c) Most individuals were found carrying one tadpole, but two and three tadpoles can also be carried at once. (d) Tadpoles transported simultaneously differ significantly in size. Photos by: B. Rojas.

### Aggressive behavior

We observed 23 agonistic encounters involving both male - male (n = 10) and female - female (n = 13) pairs. On one occasion, a male shortly attacked a female while attacking another male but resumed courting the same female shortly after. The agonistic interactions ranged from short instances of chasing without any physical contact to prolonged continuous physical combat lasting at least 20 min.

In both sexes, the physical fights involved kicking, jumping on each other’s back, and pressing either the head or the dorsum against the substrate (Fig. 9; Supplementary video 5). In most cases, we were unable to identify the origin of the conflict, but it seemed to occur both in the presence and absence of an individual of the opposite sex. While this was not always the case, both male and female aggressive interactions were observed while one of the contestants was involved in courtship. For example, on one occasion, while observing a courting pair in which the female was following the male closely, a second female who had been under a log suddenly appeared and immediately assaulted the courting female. The intruding female jumped on top of the courting female, trying to press the body of the latter against the substrate. The courting female recovered, and tried to go on top of the intruder, and these alternating attacks rapidly became a seemingly intense physical combat, in which movements and attacks occurred at a high speed. The male turned away from the females and started to call at a high repetition rate. The combat lasted for about ten minutes at the end of which the intruder female moved away, presumably defeated by the courting female. The courting pair continued to be together for a couple more hours until egg-laying occurred. On other occasions, we noticed the presence of a female in the vicinity of two males engaged in a physical combat after which one of the males courted the female while the other moved away. Some agonistic interactions both between males and between females occurred with no visible involvement of the opposite sex. Interestingly, on two occasions males carrying tadpoles where also seen engaged in physical combats with other males.

**Figure 9.**
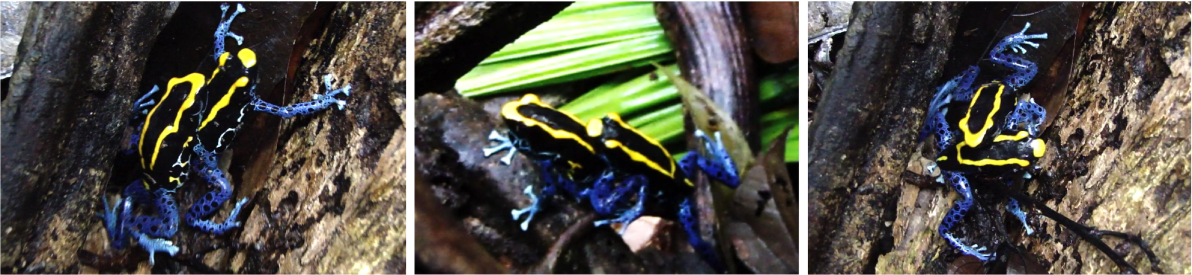
Sequence of an agonistic encounter between two male *D. tinctorius*. Physical combats involve wrestling, pressing the opponent against the substrate (with either the forelimbs or the whole body), and kicking. Occasionally males also vocalize during fighting, as seen by the inflated vocal sac in the middle photograph. Photos by: B. Rojas.

## Discussion

The purpose of this study was to provide basic information about various aspects of the natural history of *D. tinctorius* in the wild that could be used as background knowledge for future research on the behavioral ecology, evolution and conservation of the species. We describe their reproductive and social behaviors, habitat use, and their remarkable colonization of tree-fall gaps as soon as they occur. The implications of these findings, as well as some hypotheses derived from our observations, are discussed below.

*Dendrobates tinctorius* males were most often found climbing, foraging, and hiding around forest structures, such as dead logs, fallen branches, roots, tree buttresses, and palm leaves. These structures are used as oviposition sites (this study), and are also the types of structures that accumulate rainwater, forming pools where newly hatched tadpoles are deposited (Rojas 2014). Females, in contrast, were more often found foraging on the ground in open areas. Sex differences in microhabitat use might thus be related to differences in parental duties, as males periodically attend developing clutches and are in charge of tadpole transport and deposition. Forest structures can also be used by both sexes as communal retreats during dry periods (Born et al. 2010), as has also been reported for *D. truncatus*, a closely related species (Gualdrón et al. 2016). Such microhabitat likely provides higher humidity and shelter from potential predators.

Males’ association with structured habitats may also be the key reason why they are more likely to be found in tree-fall gaps long after their formation. A recent study reported higher tadpole deposition rates in pools at recent tree-fall gaps in comparison to pools in the closed forest (Rojas 2015), suggesting that the availability of new places for tadpole deposition is one of the drivers of tree-fall gap invasion in this species. However, the immediate arrival in tree-fall gaps is not exclusive of males. Females are as likely to get to new tree-fall gaps within the first three days of their formation (Fig. 4B), possibly attracted by the sudden abundance and diversity of food (pers. obs.); in fact, frogs captured in recently-formed tree-fall gaps have shown a tendency to have more prey items in their stomach than frogs caught in the closed forest (Born et al. 2010). Moreover, as suggested by Born et al. (2010), the simultaneous presence of many individuals (Fig. 4A) can make tree-fall gaps a perfect mating arena.

The mechanism by which these frogs detect and locate treefalls remains unknown. However, the sound and seismic cues produced during a treefall might be sufficient, as some frogs are known to detect vibrational signals from conspecifics (Lewis and Narins 1985; Caldwell et al. 2010), heterospecifics (Warkentin et al. 2007) and abiotic factors like rain (Caldwell et al. 2010); these three kinds of signals are presumably much weaker than those produced by a treefall. Low-frequency seismic cues could be detected at long distances but are short in duration; thus, it is possible that strong olfactory cues and light gradients produced by a fresh treefall provide the additional information needed for orientation.

One of the most unusual aspects of *D. tinctorius*’ reproductive behavior, and likely one of the reasons why their behavior has been rarely studied in the wild (but see Born et al. 2010; Rojas 2014, 2015; Rojas et al. 2014a for examples of field studies on the species), is the lack of advertisement calls. Most male frogs, including other dendrobatids, use calls to attract females and to repel rival males (Gerhardt and Huber 2002; Erdtmann and Amézquita 2009; Santos et al. 2014), making them also easier to locate by researchers. The structure of these calls shows great variation across the poison frog family (Erdtmann and Amézquita 2009), and a recent large-scale comparative study (Santos et al. 2014) argued that a reduced predation pressure has facilitated this diversification in acoustic signals in aposematic species. Paradoxically, and in contrast to the vast majority of frogs, aposematic *D. tinctorius* appears to have lost the advertisement function of its call altogether. Two closely related species, *D. auratus* and *D. truncatus*, also vocalize less frequently and at lower intensities than most other poison frogs, but still use calling both for territorial advertisement and courtship (Wells 1978; Summers 1989; Erdtmann and Amézquita 2009; Gualdrón-Duarte et al. 2016). What factors drove or facilitated the loss of typical calling behavior in *D. tinctorius* remains an intriguing evolutionary puzzle. Despite their toxicity, recent studies indicate that predation risk by naïve predators may still be an important selective pressure (Noonan and Comeault 2009; Comeault and Noonan 2011; Rojas et al. 2014b), suggesting that the increased exposure associated with prominent calling behavior should be selected against. However, this situation is not exclusive to *D. tinctorius*, and poison frogs in the genus *Oophaga*, for instance, have kept their advertisement calls and an active vocal behavior despite their conspicuous coloration (Pröhl 2003; Vargas-Salinas and Amézquita 2013; Willink et al. 2013). On the other hand, male and female *D. tinctorius* tend to segregate in and around tree-fall gaps and other forest structures, potentially facilitating mating pair formation by direct encounter without the need of acoustic signals. We speculate that such microhabitat segregation and the availability of putative visual signals for communication in a diurnal colorful frog (discussed below) has promoted the loss of the advertisement call in *D. tinctorius*.

Male *D. tinctorius* use calls, however, in courtship and agonistic interactions. The courtship calls resembles a lower intensity version of calls produced by closely related species, such as *D. auratus* and *D. truncatus* (Wells 1978, Gualdrón et al. 2016; BR, pers. obs.). In addition to advertisement calls, many other dendrobatid frogs use soft courtship calls (e.g., Roithmair 1994), which are thought to facilitate the contact with the female during the prolonged courtship while reducing the potential detection and conflict with competitor males (Wells 2007). Courtship calls may also stimulate the ovulation in females, signal territory ownership or function as visual signals because of the slow and prominent vocal sac inflation. In *D. tinctorius*, males often take a distinct elevated posture when calling both during courtship and agonistic encounters, and this posture is retained at times in the absence of vocalizations. This so-called ‘upright posture’ is thought to function as a visual signal in both contexts (Hödl and Amézquita 2001). Visual signals (Summers et al. 1999, Santos et al. 2014, Narins et al. 2003, de Luna et al. 2010) and tactile interactions (Bourne et al. 2001, Pröhl & Hödl 1999, Summers 1992) have been long thought to play an important role in poison frog communication. Aspects of dorsal coloration, for example, are known to influence mating decisions (Summers et al. 1999; Maan and Cummings 2008) and agonistic encounters (Crothers et al. 2011) in at least one species of poison frog, *O. pumilio*. However, in *O. pumilio* and other species, acoustic signals still mediate the initial mate attraction (Lötters et al. 2003; Pröhl 2003; Dreher and Pröhl 2014) and male-male competition (Crump 1972; Bee 2003; Amézquita et al. 2006; Rojas et al. 2006; Tumulty et al. 2018). In the absence of advertisement calls, the use of tactile stimuli and both static (such as dorsal colour patterns) and dynamic visual signals most likely plays a predominant role in *D. tinctorius* communication. Dorsal color patterns might mediate mate choice (Rojas 2017), given that individuals follow each other for a considerable amount of time while searching for a suitable place for oviposition. Males have been found to have a higher proportion of yellow in their dorsal area than females in our study population (Rojas and Endler 2013). This has been suggested to be particularly beneficial during tadpole transport (Rojas and Endler 2013), a task that requires long displacements and prolonged exposure, especially when climbing trees (A. Pasukonis, M. Loretto and B. Rojas, unpubl. data). Male coloration might thus indicate parental male quality and be subject to sexual selection (Rojas 2017). We suggest, however, that the variable coloration patterns on these frogs’ front, forelimbs, and flanks, also have the potential to be used as signals, as a lot of the time the frogs are either facing or next to each other during courtship (Rojas 2012). These color patterns may also be used for species, sex, or even individual recognition from the distance. Individual recognition has not been shown in any amphibian, but the relatively complex social behavior, the lack of acoustic communication, and the repeated encounters in their shared micro-habitat might have promoted such ability in *D. tinctorius*. Both male and female *D. tinctorius* engage in intra-sex aggression that may escalate to intense physical combats, which involve chasing, wrestling and prolonged pressure over the opponent’s head or dorsum. These types of behaviors have been also reported for the closely related *D. auratus* (Wells 1978; Summers 1989) and *D. leucomelas* (Summers 1992). Aggression in male poison frogs is usually a result of male competition for mates and territorial defense mediated by acoustic interactions (reviewed in Pröhl 2005). To the best of our knowledge, males of all dendrobatid species studied to date show some degree of territoriality (Pröhl 2005). *Dendrobates tinctorius* seems also unusual in this respect, as they do not appear to defend exclusive areas. Similar to Born et al. (2010), we have observed males foraging in close proximity without aggressive escalations in large aggregations around fresh tree-fall gaps, as well as around structures where a few males might take refuge. We observed that the presence of individuals of the opposite sex, especially during courtship, was the cause of some of the agonistic encounters both between males and between females. Inter-female aggression has been also reported for *Mannophryne trinitatis* (Wells 1980), *D. auratus (Wells 1978; Summers 1989), D. leucomelas (Summers 1992*), and *O. pumilio* (Meuche et al. 2011). Just like in *D. tinctorius*, in the closely related *D. auratus*, tadpoles are cannibalistic and males may deposit tadpoles from multiple clutches in the same pool (Summers 1989, 1990). As suggested for *D. auratus*, female aggression thus might be the result of attempts to monopolize males and reduce the potential competition and risk of cannibalism by unrelated tadpoles in shared pools. Interestingly, we also observed aggressive interactions that seemingly did not involve a third individual, suggesting aggression triggers other than access to mates. These observations should, however, be interpreted with caution, as we cannot be certain that a third individual was not hiding in the area. In addition to mating context, aggression in some dendrobatid frogs has been linked to defense of shelter and feeding areas (Wells 19080; Meuche et al. 2011), but *D. tinctorius* does not appear to defend exclusive territories (Born et al. 2010). Some of the aggressive interactions resulted in the defeated individual being chased away, as if in a territorial displacement, but others terminated with both individuals continuing to forage nearby. This hints at an establishment of dominance hierarchies between opponents, which we suggest could be the result of repeated encounters of individuals in their shared microhabitat. Dominance hierarchies are well documented in all other vertebrates, where dominant individuals get preferential access to food, mates, and shelter (reviewed in Huntingford 2013). However, the formation of such potential hierarchies has not been described for any anuran species in the wild, despite being suggested to arise among poison frogs in captivity (Zimmermann and Zimmermann 1988) and to mediate conflict resolution in *O. lehmanni* (Rojas 2002). This is, therefore, a subject that merits further investigation. Aggressive behavior and territoriality in *D. tinctorius* might be context-dependent and related to population density, variation in food abundance and other resources, such as structures for shelter or oviposition. In the absence of vocalizations, *D. tinctorius* may be using visual signals to get information about the fighting abilities of their opponents, as it has been reported for male *O. pumilio* (Crothers et al. 2011; Crothers and Cummings 2015), and settle their conflicts before escalating to physical combats (Rojas 2017). Social behavior in *D. tinctorius* is a promising avenue of research, which could provide insights into the evolution of visual communication and factors influencing anuran aggressive and territorial behavior in the absence of acoustic communication.

Egg clutches at our study site have high mortality and are much smaller than those reported in captivity, which may have up to 14 eggs (Lötters et al. 2007). This does not seem to be an exception, as levels of hatching failure of up to 80% have been previously reported for *O. pumilio* (Pröhl and Hödl 1999). Loss of most eggs or embryos is likely due to predation (e.g., Juncá and Rodrigues 2006), or to fungal infections (Fig 6B). On one occasion, we observed a female unrelated to the clutch on top of the missing eggs, indicating possible cannibalism. This behavior has been previously reported in *D. auratus* as a mechanism of intra-female competition (Summers 1989; 1990).

Upon hatching, males take tadpoles, either all of them or one at a time, to bodies of water. The latter is thought to be the case of most *Dendrobates*, although the evidence supporting this pattern comes mostly from observations in captivity (Lötters et al. 2007). Transport of single tadpoles, one by one, implies several trips between the place where clutch was laid and the pools, a task that has been shown to require remarkable spatial abilities (Pašukonis et al. 2016) and probably a high energetic cost (Beck et al. 2017). In *D. tinctorius*, males carry one or two (sometimes three) tadpoles at a time. In combination with the high clutch mortality rates observed, this suggests that males often take all the larvae that survive within a given clutch at once. However, we have observed at least a few instances in which males take one tadpole to a pool and then return to get the rest (A. Pašukonis, M. Loretto and B. Rojas, unpubl. data). Furthermore, males have been seen depositing one tadpole in the same pool where they had deposited another tadpole (Rojas 2014), which could also mean that males carrying two tadpoles at once are transporting tadpoles from different clutches. This is supported by the fact that tadpoles transported at the same time on a male’s back differ significantly in size. In *D. auratus* for example, males have been seen moistening a fresh clutch and a hatching clutch within the same hour in captivity (Wells 1978), and attending multiple clutches of different stages in the field (Wells 1978; Summers 1990). However, these differences in size could also be inherent to within-clutch variation in hatching size. Size difference between tadpoles transported simultaneously was particularly noticeable in the tadpoles on the back of the only female found performing these duties. While rare (1 in >100 tadpole transport events reported here), tadpole transport might be taken over by females if males go missing. This type of flexible compensation is known to occur in at least one other species of poison frog, *Allobates femoralis* (Ringler et al. 2015). As reported before, we observed tadpole deposition in different water-holding structures in the forest, from palm bracts on the ground to tree holes high up. However, the specific characteristics that influence pool choice by a male and favor successful tadpole development are currently unknown. It has been previously suggested that, despite the high levels of tadpole cannibalism, parents might use the presence of larger tadpoles as a cue of pool quality. Presence of large tadpoles may indicate that basic requirements, such as sufficient nutrients and water stability, have been met to allow tadpole development (Rojas 2014). Even less is understood about the role that *D. tinctorius* plays in the ecology of other phytotelm-breeding anurans, especially considering that most species are restricted to terrestrial or arboreal habitat; meanwhile, *D. tinctorius* and their carnivorous tadpoles are capable of exploiting pools at all heights (Gaucher 2002). How *D. tinctorius* finds canopy pools is unknown, but it has been speculated that they may eavesdrop on the calls of treefrog species such as *Trachycephalus resinifictrix* and *T. hadroceps*, which breed in arboreal water bodies (Gaucher 2002). We further hypothesize that enlarged male toe-pads (apt for climbing) and aposematic coloration gave *D. tinctorius* access to a wider variety of aquatic habitats despite being exposed to would-be predators for prolonged periods of time during tadpole transport.

Approximately 43% of the amphibian species worldwide are experiencing population declines (Stuart, et al. 2004), largely as a consequence of the spread of a deadly disease caused by the fungus *Batrachochytrium dendrobatidis (Bd*) (Lips et al. 2006; Lötters et al. 2009; Lips 2016; Bower et al. 2017). Despite having a low *Bd* prevalence compared to species in other families and regions (e.g. Flechas et al. 2012), phytotelm-breeding dendrobatids, including *D. tinctorius*, have been found to have the highest prevalence of *Bd* in recent studies done in French Guiana, indicating that dendrobatid species may be more vulnerable to *Bd* infection than previously suspected (Courtois et al. 2015). Furthermore, the same study reports an increase in *Bd* prevalence in *D. tinctorius* between 2009 and 2012, hinting at a relatively recent *Bd* establishment and its current spread in French Guiana. While *Bd* research has been mostly focused on adult frogs, it is known that tadpoles can also get infected due to their keratinized mouthparts (Berger et al. 1998; Blaustein et al. 2005). However, there is currently no information on *Bd* prevalence in *D. tinctorius* tadpoles, or studies assessing the presence of *Bd* in the pools where tadpoles develop, despite reports of *Bd* occurrence in phytotelmata and phytotelm-breeding frog species in other Neotropical areas such as Panama and Ecuador (Cossel and Lindquist 2009; McCracken et al. 2009). Furthermore, considering that dispersal of *Bd* outside large bodies of water may imply an amphibian vector (Kolby et al. 2015), we urge studies evaluating the role of *D. tinctorius* adults as *Bd* vectors across forest strata (i.e., from the ground to the canopy), and advocate that increasing our knowledge on the behavior and ecology of this species may improve our understanding on the dynamics of *Bd* transmission in their habitat.

Another major threat for anurans is habitat destruction (Cushman 2006). While our study population occurs within in a natural reserve, many populations of *D. tinctorius* are in unprotected areas, which are under threat primarily by gold mining-driven deforestation. It is estimated that approximately 41% (~684 km2) of the deforestation in the South American tropical rainforest between 2001 and 2013 occurred in the so-called Guianan moist forest ecoregion due to gold mining activities (Alvarez-Berríos and Aide 2015). Because *D. tinctorius* is often distributed in small patchy populations (Noonan and Gaucher 2006), deforestation even at the small-scales used for gold mining, can have a detrimental, probably irreversible effect on the life histories and survival of this species and other phytotelm-breeders. We thus fully support the long-term monitoring strategies suggested by Courtois et al. (2015) and currently implemented across several nature reserves in French Guiana (e.g., http://www.reserve-tresor.fr/en/our-actions/studies-and-surveys/herpetology) to allow the timely assessment of changes in population size and sudden declines, especially of ‘sentinel species’ such as *D. tinctorius* (Courtois et al. 2013, 2015). These types of strategies, together with basic research on the natural history of threatened species are key, not only for the formulation of successful conservation policies, but also for the education and future engagement of public essential for the preservation of wildlife at a local scale.

## Additional Information and Declarations

### Author Contributions

Both authors collected information in the field, analyzed data, and prepared figures for the paper. BR wrote a first draft of the manuscript, which was then edited and approved by AP.

### Animal Ethics

Our research was approved and authorized by the scientific committee of the Nouragues Ecological Research Station. We strictly adhered to the current French and European Union law, and followed the ‘Guidelines for use of live amphibians and reptiles in the field and laboratory research’ by the Herpetological Animal Care and Use Committee (HACC) of the American Society of Ichthyologists and Herpetologists and the ‘Guidelines for the use of animals in teaching and research’ by the Association for the Study of Animal Behaviour (ASAB).

## Supplementary videos

1. Tadpole transport and pool inspection https://youtu.be/KE5LB-2IsTU

2. Cannibalism/tadpole aggression https://youtu.be/Utvrnqi-VOk

3. Courtship https://youtu.be/gNlZLBpivMI

4. Egg laying https://youtu.be/zf-4aOXeir0

5. Male and female aggressive behavior https://www.youtube.com/watch?v=g5x_K0x7Lcg&feature=youtu.be

## Acknowledgements

We are extremely grateful to John A. Endler and Walter Hödl for insightful and encouraging discussions on the importance of natural history studies. Jennifer Devillechabrolle, Diana Pizano, Oscar Ramos, Valentine Alt, and Matthias-Claudio Loretto provided invaluable field assistance. Philippe Gaucher, Mathias Fernandez, Gilles Peroz and the rest of Les Nouragues Ecological Research station staff helped with logistics; P. G. also shared helpful information on many years observing these frogs in the wild. We are thankful to Eva Fischer for providing the excellent photo of the tadpole’s mouthparts, Antoine Fouquet for providing photos of two populations of *D. tinctorius*, and Matthias-Claudio Loretto for courtship videos.

## References

Alvarez-Berríos, N. L., and T. M. Aide. 2015. Global demand for gold is another threat for tropical forests. Environmental research letters: ERL [Web site] 10:014006.

Amézquita, A., L. Castellanos, and W. Hödl. 2005. Auditory matching of male Epipedobates femoralis (Anura: Dendrobatidae) under field conditions. Animal behaviour 70:1377–1386.

Amézquita, A., S. V. Flechas, A. P. Lima, H. Gasser, and W. Hödl. 2011. Acoustic interference and recognition space within a complex assemblage of dendrobatid frogs. Proceedings of the National Academy of Sciences of the United States of America 108:17058–17063.

Amézquita, A., W. Hödl, A. P. Lima, L. Castellanos, L. Erdtmann, and M. C. de Araújo. 2006. Masking interference and the evolution of the acoustic communication system in the Amazonian dendrobatid frog Allobates femoralis. Evolution; international journal of organic evolution 60:1874–1887.

Barnett, J. B., C. Michalis, N. E. Scott-Samuel, and I. C. Cuthill. 2018. Distance-dependent defensive coloration in the poison frog Dendrobates tinctorius, Dendrobatidae. Proceedings of the National Academy of Sciences of the United States of America 115:6416–6421.

Beck, K. B., M.-C. Loretto, M. Ringler, W. Hödl, and A. Pašukonis. 2017. Relying on known or exploring for new? Movement patterns and reproductive resource use in a tadpole-transporting frog. PeerJ 5:e3745.

Beeby, A. 2001. What do sentinels stand for? Environmental pollution 112:285–298.

Bee, M. A. 2003. A test of the “dear enemy effect” in the strawberry dart-poison frog (Dendrobates pumilio). Behavioral ecology and sociobiology.

Berger, L., R. Speare, P. Daszak, D. E. Green, A. A. Cunningham, C. L. Goggin, R. Slocombe, et al. 1998. Chytridiomycosis causes amphibian mortality associated with population declines in the rain forests of Australia and Central America. Proceedings of the National Academy of Sciences of the United States of America 95:9031–9036.

Blaustein, A. R., J. M. Romansic, E. A. Scheessele, B. A. Han, A. P. Pessier, and J. E. Longcore. 2005. Interspecific Variation in Susceptibility of Frog Tadpoles to the Pathogenic Fungus Batrachochytrium dendrobatidis. Conservation biology: the journal of the Society for Conservation Biology 19:1460–1468.

Boersma, P., and D. Weenink. 2014. Praat: Doing Phonetics by Computer [Computer software]. Version 5.3. 84.

Born, M., F. Bongers, E. H. Poelman, and F. J. Sterck. 2010. Dry-season retreat and dietary shift of the dart-poison frog Dendrobates tinctorius (Anura: Dendrobatidae). Phyllomedusa: journal of neotropical herpetology 9:37–52.

Bower, D. S., K. R. Lips, L. Schwarzkopf, A. Georges, and S. Clulow. 2017. Amphibians on the brink. Science 357:454–455.

Brodie, E. D., and M. S. Tumbarello. 1978. The antipredator functions of Dendrobates auratus (Amphibia, Anura, Dendrobatidae) skin secretion in regard to a snake predator (Thamnophis). Journal of herpetology 12:264–265.

Brown, J. L., V. Morales, and K. Summers. 2010. A key ecological trait drove the evolution of biparental care and monogamy in an amphibian. The American naturalist 175:436–446.

Brown, J. L., E. Twomey, A. Amezquita, M. B. de Souza, J. P. Caldwell, S. Loetters, R. May, et al. 2011. A taxonomic revision of the Neotropical poison frog genus Ranitomeya (Amphibia: Dendrobatidae). Zootaxa 3083:1–120.

Caldwell, J. P. 1997. Pair bonding in spotted poison frogs. Nature 385:211.

Caldwell, M. S., G. R. Johnston, J. G. McDaniel, and K. M. Warkentin. 2010. Vibrational signaling in the agonistic interactions of red-eyed treefrogs. Current biology: CB 20:1012–1017.

Comeault, A. A., and B. P. Noonan. 2011. Spatial variation in the fitness of divergent aposematic phenotypes of the poison frog, Dendrobates tinctorius. Journal of evolutionary biology 24:1374–1379.

Cossel, J. O., Jr, and E. D. Lindquist. 2009. AMPHIBIAN DISEASES-Batrachochytrium dendrobatidis in Arboreal and Lotic Water Sources in Panama. Herpetological review.

Courtois, E. A., J. Devillechabrolle, M. Dewynter, K. Pineau, P. Gaucher, and J. Chave. 2013. Monitoring strategy for eight amphibian species in French Guiana, South America. PloS one 8:e67486.

Courtois, E. A., P. Gaucher, J. Chave, and D. S. Schmeller. 2015. Widespread occurrence of bd in French Guiana, South America. PloS one 10:e0125128.

Courtois, E. A., K. Pineau, B. Villette, D. S. Schmeller, and P. Gaucher. 2012. Population estimates of Dendrobates tinctorius (Anura: Dendrobatidae) at three sites in French Guiana and first record of chytrid infection. Phyllomedusa: journal of neotropical herpetology 11:63–70.

Crothers, L., E. Gering, and M. Cummings. 2011. Aposematic signal variation predicts male–male interactions in a polymorphic poison frog. Evolution; international journal of organic evolution.

Crothers, L. R., and M. E. Cummings. 2015. A multifunctional warning signal behaves as an agonistic status signal in a poison frog. Behavioral ecology: official journal of the International Society for Behavioral Ecology 26:560–568.

Crump, M. L. 1972. Territoriality and Mating Behavior in Dendrobates granuliferus (Anura: Dendrobatidae). Herpetologica 28:195–198.

Cushman, S. A. 2006. Effects of habitat loss and fragmentation on amphibians: A review and prospectus. Biological conservation 128:231–240.

Daly, J. W., C. W. Myers, and N. Whittaker. 1987. Further classification of skin alkaloids from neotropical poison frogs (Dendrobatidae), with a general survey of toxic/noxious substances in the amphibia. Toxicon: official journal of the International Society on Toxinology 25:1023–1095.

Donnelly, M. A. 1989. Effects of reproductive resource supplementation on space-use patterns in Dendrobates pumilio. Oecologia 81:212–218.

Dreher, C. E., and H. Pröhl. 2014. Multiple sexual signals: calls over colors for mate attraction in an aposematic, color-diverse poison frog. Frontiers in Ecology and Evolution 2:3224.

Endler, J. A. 2015. Writing scientific papers, with special reference to Evolutionary Ecology. Evolutionary ecology 29:465–478.

Erdtmann, L., and A. Amézquita. 2009. Differential Evolution of Advertisement Call Traits in Dart-Poison Frogs (Anura: Dendrobatidae). Ethology: formerly Zeitschrift fur Tierpsychologie 115:801–811.

Fandiño, M. C., H. Lüddecke, and A. Amézquita. 1997. Vocalisation and larval transportation of male Colostethus subpunctatus (Anura: Dendrobatidae). Amphibia-reptilia: publication of the Societas Europaea Herpetologica 18:39–48.

Flechas, S. V., C. Sarmiento, and A. Amezquita. 2012. Bd on the Beach: high prevalence of *Batrachochytrium dendrobatidis* in the lowland forests of Gorgona island (Colombia, South America). Ecohealth 9:298–302.

Forsman, A., and M. Hagman. 2006. Calling is an honest indicator of paternal genetic quality in poison frogs. Evolution; international journal of organic evolution 60:2148–2157.

Gaucher, P. 2002. Premières données sur Phrynohyas hadroceps, Rainette arboricole du plateau des Guyanes (Amphibia:Anura:Hylidae) (Révision taxonomique, éco-éthologie de la reproduction).

Gerhardt, C. H., and F. Huber. 2002. Acoustic Communication in Insects and Anurans: Common Problems and Diverse Solutions. University of Chicago Press.

Gorzula, S. 1996. The Trade in Dendrobatid Frogs. Herpetological review 27:3.

Grant, T., D. R. Frost, J. P. Caldwell, R. Gagliardo, C. F. B. Haddad, P. J. R. Kok, D. B. Means, et al. 2006. Phylogenetic systematics of dart-poison frogs and their relatives (Amphibia: Athesphatanura: Dendrobatidae). Bulletin of the American Museum of Natural History 1–262.

Gualdrón-Duarte, J. E., V. F. Luna-Mora, and Rivera-Correa M Kahn T. 2016. Yellow-striped Poison Frog Dendrobates truncatus (Cope, 1861 “1860”). Pages 323–328 in T. R. Kahn, E. La Marca, S. Lötters, J. L. Brown, E. Towney, and A. Amézquita, eds. Aposematic Poison Frogs (Dendrobatidae) of the Andean Countries: Bolivia, Colombia, Ecuador, Perú and Venezuela, Conservation International Tropical Field Guides Series. Conservation International, Arlignton. USA.

Hödl, W., and A. Amézquita. 2001. Visual signaling in anuran amphibians. Anuran communication 121–141.

Hödl, W., A. Amézquita, and P. M. Narins. 2004. The role of call frequency and the auditory papillae in phonotactic behavior in male Dart-poison frogs Epipedobates femoralis (Dendrobatidae). Journal of comparative physiology. A, Neuroethology, sensory, neural, and behavioral physiology 190:823–829.

Hoogmoed, M., and Avila-Pires, T. C. S. 2012. Inventory of color polymorphism in populations of *Dendrobates galactonotus* (Anura: Dendrobatidae), a poison frog endemic to Brazil. Phyllomedusa 11:95–115.

Huntingford, F. A. 2013. Animal Conflict. Springer Science & Business Media.

Juncá, F. A., and M. T. Rodrigues. 2006. The reproductive success of Colostethus stepheni(Anura: Dendrobatidae). Studies on neotropical fauna and environment 41:9–17.

Kolby, J. E., S. D. Ramirez, L. Berger, K. L. Richards-Hrdlicka, M. Jocque, and L. F. Skerratt. 2015. Terrestrial Dispersal and Potential Environmental Transmission of the Amphibian Chytrid Fungus (Batrachochytrium dendrobatidis). PloS one 10:e0125386.

Lewis, E. R., and P. M. Narins. 1985. Do frogs communicate with seismic signals? Science 227:187–189.

Lips, K. R. 2016. Overview of chytrid emergence and impacts on amphibians. Philosophical transactions of the Royal Society of London. Series B, Biological sciences 371.

Lips, K. R., F. Brem, R. Brenes, J. D. Reeve, R. A. Alford, J. Voyles, C. Carey, et al. 2006. Emerging infectious disease and the loss of biodiversity in a Neotropical amphibian community. Proceedings of the National Academy of Sciences of the United States of America 103:3165–3170.

Longcore, J. E., A. P. Pessier, and D. K. Nichols. 1999. *Batrachochytrium Dendrobatidis gen. et sp. nov*., a Chytrid Pathogenic to Amphibians. Mycologia 91:219–227.

Lötters, S., K.-H. Jungfer, F. W. Henkel, and W. Schmidt. 2007. Poison Frogs: Biology, Species & Captive Care. Edition Chimaira.

Lötters, S., J. Kielgast, J. Bielby, S. Schmidtlein, J. Bosch, M. Veith, S. F. Walker, et al. 2009. The link between rapid enigmatic amphibian decline and the globally emerging chytrid fungus. EcoHealth 6:358–372.

Lötters, S., S. Reichle, and K.-H. Jungfer. 2003. Advertisement calls of Neotropical poison frogs (Amphibia: Dendrobatidae) of the genera Colostethus, Dendrobates and Epipedobates, with notes on dendrobatid call classification. Journal of natural history 37:1899–1911.

Maan, M. E., and M. E. Cummings. 2008. Female preferences for aposematic signal components in a polymorphic poison frog. Evolution; international journal of organic evolution 62:2334–2345.

Maan, M. E., and M. E. Cummings. 2009. Sexual dimorphism and directional sexual selection on aposematic signals in a poison frog. Proceedings of the National Academy of Sciences of the United States of America 106:19072–19077.

McCracken, S., J. P. Gaertner, and M. R. J. Forstner. 2009. Detection of Batrachochytrium dendrobatidis in amphibians from the forest floor to the upper canopy of an Ecuadorian Amazon lowland rainforest. Herpetological.

Meuche, I., K. E. Linsenmair, and H. Pröhl. 2011. Female Territoriality in the Strawberry Poison Frog (Oophaga pumilio). Copeia 2011:351–356.

Myers, C. W., and J. W. Daly. 1976. Preliminary evaluation of skin toxins and vocalizations in taxonomic and evolutionary studies of poison-dart frogs (Dendrobatidae). Bulletin of the AMNH; v. 157, article 3.

Myers, C. W., and J. W. Daly. 1980. Taxonomy and ecology of Dendrobates bombetes, a new Andean poison frog with new skin toxins. American Museum novitates; no. 2692.

Myers, C. W., and J. W. Daly. 1983. Dart-poison frogs. Scientific American 248:96–105.

Myers, C. W., J. W. Daly, and B. Malkin. 1978. A dangerously toxic new frog (*Phyllobates*) used by Emberá indians of western Colombia, with discussion of blowgun fabrication and dart poisoning. Bull. Amer. Mus. Nat. Hist. 161:309–365.

Nijman, V., and C. R. Shepherd. 2010. The role of Asia in the global trade in CITES II-listed poison arrow frogs: hopping from Kazakhstan to Lebanon to Thailand and beyond. Biodiversity and conservation 19:1963–1970.

Noonan, B. P., and A. A. Comeault. 2009. The role of predator selection on polymorphic aposematic poison frogs. Biology letters 5:51–54.

Noonan, B. P., and P. Gaucher. 2006. Refugial isolation and secondary contact in the dyeing poison frog Dendrobates tinctorius. Molecular ecology 15:4425–4435.

Pašukonis, A., K. Trenkwalder, M. Ringler, E. Ringler, R. Mangione, J. Steininger, I. Warrington, et al. 2016. The significance of spatial memory for water finding in a tadpole-transporting frog. Animal behaviour 116:89–98.

Pašukonis, A., I. Warrington, M. Ringler, and W. Hödl. 2014. Poison frogs rely on experience to find the way home in the rainforest. Biology letters 10:20140642.

Pröhl, H. 2002. Population differences in female resource abundance, adult sex ratio, and male mating success in Dendrobates pumilio. Behavioral ecology: official journal of the International Society for Behavioral Ecology 13:175–181.

Pröhl, H. 2003. Variation in male calling behaviour and relation to male mating success in the strawberry poison frog (*Dendrobates pumilio*). Ethology 109:273–290.

Pröhl, H. 2005. Territorial Behavior in Dendrobatid Frogs. Journal of herpetology 39:354–365.

Pröhl, H., and W. Hödl. 1999. Parental investment, potential reproductive rates, and mating system in the strawberry dart-poison frog, Dendrobates pumilio. Behavioral ecology and sociobiology 46:215–220.

Ringler, E., A. Pašukonis, W. T. Fitch, L. Huber, W. Hödl, and M. Ringler. 2015. Flexible compensation of uniparental care: female poison frogs take over when males disappear. Behavioral ecology 26:1219–1225.

Roithmair, M. E. 1994. Field studies on reproductive behaviour in two dart-poison frog species (Epipedobates femoralis, Epipedobates trivittatus) in Amazonian Peru. Herpetological Journal 4:77–85.

Rojas, B. 2002. Intrinsic determinants of the outcome of agonistic encounters in the poison-arrow frog Dendrobates lehmanni (Anura: Dendrobatidae). B.Sc. Thesis, University of Los Andes, Bogota, Colombia, 36 p.

Rojas, B. 2014. Strange parental decisions: fathers of the dyeing poison frog deposit their tadpoles in pools occupied by large cannibals. Behavioral Ecology and Sociobiology 68:551–559.

Rojas, B. 2014. 2015. Mind the gap: treefalls as drivers of parental tradeoffs. Ecology and evolution 5:4028–4036.

Rojas, B. 2017. Behavioural, ecological, and evolutionary aspects of diversity in frog colour patterns. Biological Reviews 92:1059–1080.

Rojas, B., A. Amézquita, and A. Delgadillo. 2006. Matching and symmetry in the frequency recognition curve of the poison frog *Epipedobates trivittatus*. Ethology 112:564–571.

Rojas, B., J. Devillechabrolle, and J. A. Endler. 2014a. Paradox lost: variable colour-pattern geometry is associated with differences in movement in aposematic frogs. Biology Letters 10:20140193.

Rojas, B., and J. A. Endler. 2013. Sexual dimorphism and intra-populational colour pattern variation in the aposematic frog *Dendrobates tinctorius*. Evolutionary Ecology 27:739–753.

Rojas, B., P. Rautiala, and J. Mappes. 2014b. Differential detectability of polymorphic warning signals under varying light environments. Behavioural Processes 109 B:164–172.

RStudio Team. 2015. RStudio: integrated development for R.

Santos, J. C., M. Baquero, C. Barrio-Amorós, L. A. Coloma, L. K. Erdtmann, A. P. Lima, and D. C. Cannatella. 2014. Aposematism increases acoustic diversification and speciation in poison frogs. Proceedings of the Royal Society of London B 281:20141761.

Santos, J. C., L. A. Coloma, and D. C. Cannatella. 2003. Multiple, recurring origins of aposematism and diet specialization in poison frogs. Proceedings of the National Academy of Sciences 100:12792–12797.

Schmidt, W., and F. W. Henkel. 1995. Pfeilgiftfrösche im Terrarium. Landbuch-Verlag.

Schneider, C. A., W. S. Rasband, and K. W. Eliceiri. 2012. NIH Image to ImageJ: 25 years of image analysis. Nature Methods 9:671–675.

Silverstone, P. A. 1973. Observations on the Behavior and Ecology of a Colombian Poison-Arrow Frog, the Kõkoé-Pá (Dendrobates histrionicus Berthold). Herpetologica 29:295–301.

Silverstone, P. A. 1975. A revision of the poison-arrow frogs of the genus Dendrobates Wagler. Revisión de las ranas venenosas del género Dendrobates Wagler. Natural history 21:1–55.

Summers, K. 1989. Sexual selection and intra-femalecompetition in the green poison-dart frog, Dendrobates auratus. Animal behaviour 37:797–805.

Summers, K.. 1990. Paternal care and the cost of polygyny in the green dart-poison frog. Behavioral Ecology and Sociobiology 27:307–313.

Summers, K.. 1992. Mating strategies in two species of dart-poison frogs: a comparative study. Animal behaviour 43:907–919.

Summers, K., R. Symula, M. Clough, and T. Cronin. 1999. Visual mate choice in poison frogs. Proceedings. Biological sciences / The Royal Society 266:2141–2145.

Tarvin, R. D., C. M. Borghese, W. Sachs, J. C. Santos, Y. Lu, L. A. O’Connell, D. C. Cannatella, et al. 2017. Interacting amino acid replacements allow poison frogs to evolve epibatidine resistance. Science 357:1261–1266.

Team, R. C. 2014. R: A language and environment for statistical computing. Vienna, Austria: R Foundation for Statistical Computing; 2014.

Tewksbury, J. J., J. G. T. Anderson, J. D. Bakker, T. J. Billo, P. W. Dunwiddie, M. J. Groom, S. E. Hampton, et al. 2014. Natural History’s Place in Science and Society. Bioscience 64:300–310.

Tumulty, J. P., A. Pašukonis, M. Ringler, J. D. Forester, W. Hödl, and M. A. Bee. 2018. Brilliant-thighed poison frogs do not use acoustic identity information to treat territorial neighbours as dear enemies. Animal behaviour 141:203–220.

Vargas-Salinas, F., and A. Amézquita. 2013. Stream noise, hybridization, and uncoupled evolution of call traits in two lineages of poison frogs: Oophaga histrionica and Oophaga lehmanni. PloS one 8:e77545.

Warkentin, K. M., M. S. Caldwell, T. D. Siok, A. T. D’Amato, and J. G. McDaniel. 2007. Flexible information sampling in vibrational assessment of predation risk by red-eyed treefrog embryos. The Journal of experimental biology 210:614–619.

Wells, K. D. 1978. Courtship and Parental Behavior in a Panamanian Poison-Arrow Frog (Dendrobates auratus). Herpetologica 34:148–155.

Wells, K. D. 1980. Social Behavior and Communication of a Dendrobatid Frog (Colostethus trinitatis). Herpetologica 36:189–199.

Wells, K. D. 2007. The Ecology and Behavior of Amphibians. University of Chicago Press.

Willink, B., E. Brenes-Mora, F. Bolaños, and H. Pröhl. 2013. Not everything is black and white: color and behavioral variation reveal a continuum between cryptic and aposematic strategies in a polymorphic poison frog. Evolution; international journal of organic evolution 67:2783–2794.

Wollenberg, K. C., S. Lötters, C. Mora-Ferrer, and M. Veith. 2008. Disentangling composite colour patterns in a poison frog species. Biological journal of the Linnean Society. Linnean Society of London 93:433–444.

Wollenberg, K. C., M. Veith, B. P. Noonan, S. Lötters, and J. M. Quattro. 2006. Polymorphism Versus Species Richness—systematics of Large Dendrobates from the Eastern Guiana Shield (Amphibia: Dendrobatidae). Copeia 2006:623–629.

Zimmermann, H., and E. Zimmermann. 1988. Etho-Taxonomie und zoogeographische artengruppenbildung bei pfeilgiftfröschen (Anura: Dendrobatidae). Salamandra 24:125–160.

